# A novel frameshift mutation in Phosphoinositide 3-kinase regulatory subunit 1 (*PIK3R1*) causes immunodeficiency and Amyotrophic Lateral Sclerosis (ALS)

**DOI:** 10.1101/2025.05.23.655625

**Authors:** Brice Calco, Colin L. Sweeney, Joseph Steiner, Tongguang Wang, Tovah E. Markowitz, Subrata Paul, Bianca Palicha, Sara Dinges, Kelsey Yoo, Lisa Henderson, Valerie McDonald, Suk See De Ravin, Benjamin Greenberg, Christa S. Zerbe, Luigi D. Notarangelo, Steven M. Holland, Avindra Nath, Farinaz Safavi

## Abstract

Mutations in *PIK3R1*, a regulatory subunit of Class I PI3K, are implicated in immune disorders and neurological conditions. We identified a novel heterozygous pathogenic frameshift mutation (c.1710dup) in *PIK3R1* in a patient with common variable immunodeficiency who developed slowly progressive Amyotrophic Lateral Sclerosis. Induced pluripotent stem cells (iPSCs) and iPSC-derived motor neurons (iMNs) demonstrated that this mutation resulted in *PIK3R1* haploinsufficiency, with downstream activation of AKT, disruption of neuronal electrical function and increased apoptosis in iPSC-derived motor neurons. Single-cell RNA sequencing (scRNA-seq) and pathway analysis of differentially expressed genes showed apoptosis pathways were upregulated in neuronal clusters from iMNs harboring the *PIK3R1*c.1710 dup mutation. Mutated iPSC-derived brain organoids were smaller than matched controls. scRNA-seq of brain organoids showed more active apoptosis in neuronal clusters of patient-derived brain organoids. These findings identify a critical and novel role for *PIK3R1* haploinsufficiency in neuronal function and survival.

## Introduction

Class I phosphoinositide 3-kinase (PI3K) is an evolutionarily conserved enzyme that regulates proliferation, differentiation, and apoptosis. The PI3K complex consists of a catalytic subunit (p110) with four isoforms (α, β, γ, δ) and a regulatory subunit, *PIK3R1* (p85α) [1]. Pathogenic mutations in *PIK3R1* affect different protein domains and are associated with various disorders, including activated PI3K-δ syndrome 2 (APDS2), characterized by hypogammaglobulinemia and immune dysregulation [2], and SHORT syndrome, marked by short stature and endocrine abnormalities [3].

Mutations in *PIK3R1* that cause APDS2 result in the production of a mutant p85α protein, which alters the binding dynamics of the p85α-p110δ complex, preventing the inhibition of p110δ catalytic activity in immune cells [4]. In contrast, *PIK3R1* mutations associated with SHORT syndrome often result in de novo truncations of p85α, leading to reduced activation of the PI3K-AKT-mTOR pathway, which in turn causes developmental delays, lipodystrophy, and distinct facial features [5].

Neurological symptoms linked to *PIK3R1* mutations have also been reported, including developmental delays, seizures, and, more recently, secondary microcephaly [6]. Despite these associations, the specific effects of *PIK3R1* mutations on neuronal function and glial cells remain unclear. Given the essential role of the PI3K pathway in neuronal and glial function, monogenic defects in this pathway might enhance our understanding of neuronal development, function, and survival, facilitating targeted treatments.

We identified a novel heterozygous frameshift mutation (c.1710dup) in *PIK3R1* in a patient with common variable immunodeficiency (CVID) who later developed slowly progressive Amyotrophic Lateral Sclerosis (ALS). Using induced pluripotent stem cell (iPSC) technology, we investigated the underlying mechanisms of this novel mutation and its downstream signaling pathway. We also evaluated the direct effects of this *PIK3R1c.1710dup* mutation and the consequent haploinsufficiency on motor neurons and brain organoids. *PIK3R1* haploinsufficiency activates the AKT pathway and affects neuronal function and survival, confirming *PIK3R1’s* novel and critical roles in the nervous system.

## Results

### Patient clinical presentation

This 44-year-old man first presented at 12 years with recurrent sinopulmonary infections, gastroenteritis, diarrhea, and human papilloma virus skin infections with hypogammaglobulinemia. He received intermittent intravenous immunoglobulin (IVIG) with improvement of IgG serum levels and clinical response. At 30 years, he developed progressive weakness, stiffness in both legs, and frequent falls. Magnetic resonance imaging (MRI) of the brain, cerebrospinal fluid evaluation, positron emission tomography, and electromyography/nerve conduction studies (EMG/NCS) were unhelpful. He was thought to have an immune-mediated myelopathy, which was treated with plasmapheresis, cyclophosphamide, and IVIG without neurologic improvement (Figure 1A).

**Figure 1.**
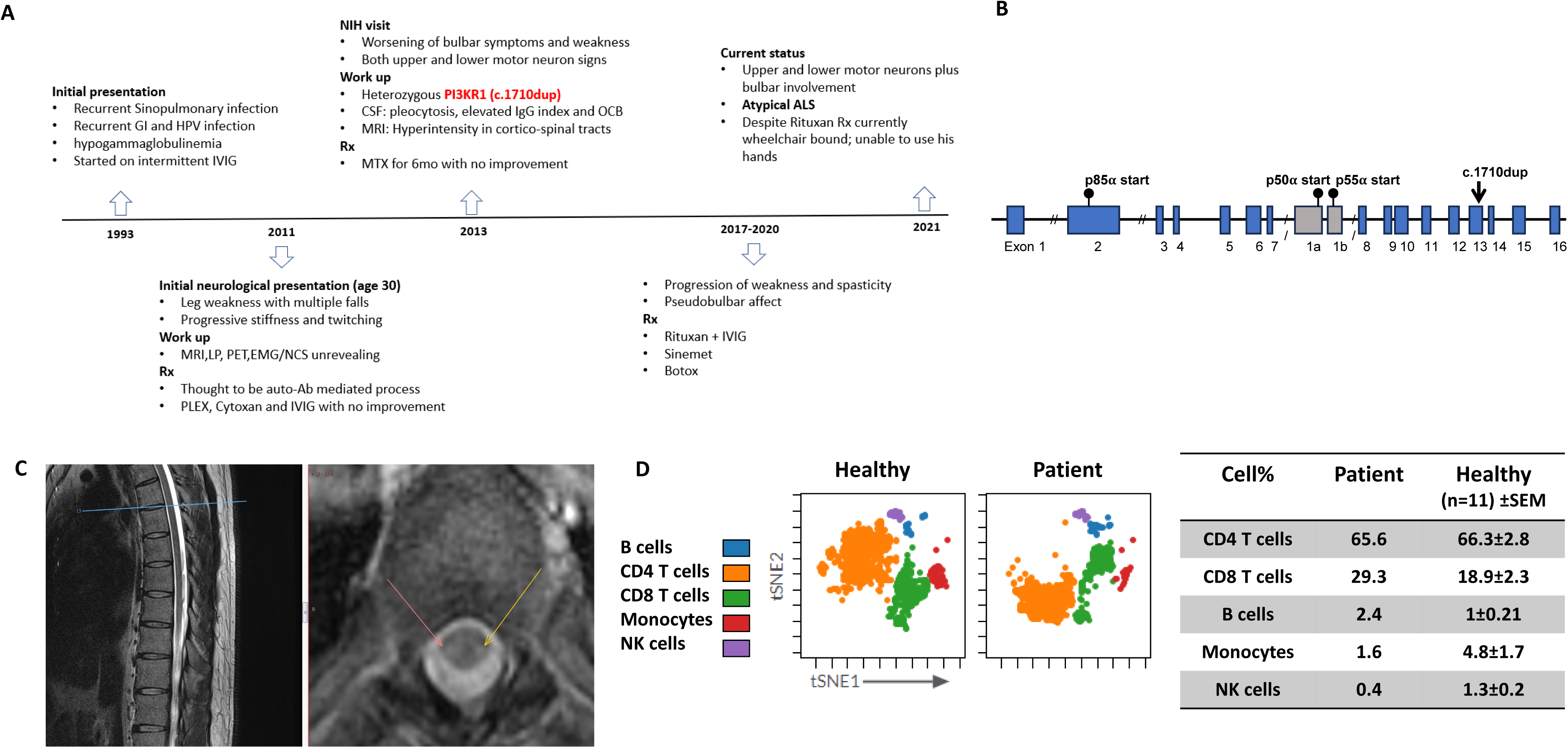
Clinical, genetic and imaging findings in patient with *PIK3Rc.11710 dup* mutation; **A)** Clinical timeline summarizing the patient’s clinical course, including the onset of symptoms concerning for immunodeficiency, neurological symptoms, work up and treatment interventions. **B)** Whole-genome sequencing (WGS) identified a novel pathogenic frameshift mutation in *PIK3R1* c.1710dup, shown in a schematic representation. **C)** MRI imaging demonstrated corticospinal tract hyperintensities in axial cuts of the spinal cord shown by arrows. **D)** CSF cells immunophenotype from healthy donors (n=11) and patient was assessed by flow cytometry and analyzed by Cytobank software. Main immune cell cluster percentages (CD4+ T cells, CD8+ T cells, B cells, Monocytes and NK cells) in CSF from patient and healthy donors (mean values +/− SEM) are shown indicating an elevated CD8+ T cell and B cell percentages in patient’s CSF compared to healthy donors.

On evaluation at 32 years, he had IgG 604 [642-1730 mg/dl], IgM <6 [34-342 mg/dl], IgA <8 [91-499 mg/dl] despite normal numbers of T and B cells (Supplemental Figure 1A). Bone marrow biopsy was hypocellular with adequate early B cells, markedly decreased mature B cells, and absent plasma cells. Whole-genome sequencing (WGS) identified a novel heterozygous frameshift variant in *PIK3R1* (c.1710dup) (Figure 1B).predicted to result in early truncation of all three isoforms of the encoded protein (p.I571Yfs*31 for the p85α isoform, p.I301Yfs*31 for the p55α isoform, and p.I271Yfs*31 for the p50α isoform).

The patient developed progressive worsening of motor function, leg stiffness, spasticity and bulbar symptoms, included dysphagia and dysarthria with preserved cognition. Cerebrospinal fluid (CSF) analyses showed intermittent lymphocytic pleocytosis, elevated IgG, and isolated oligoclonal bands (OCBs) (Supplemental Figure 1B). Spinal cord MRI showed hyperintensities in the corticospinal tracts (Figure 1C). Serum and CSF autoimmune encephalopathy and neuropathy panels were negative. EMG/NCS showed active denervation in motor neurons with increased amplitude of the motor units, reduced recruitment positive waves, and fasciculations and fibrillations in several muscle groups in upper and lower limbs, all consistent with motor neuron disease. Based on the CSF pleocytosis and elevated CSF IgG and demonstration of a *PIK3R1* mutation, which is frequently associated with immune dysregulation, methotrexate (MTX) and sirolimus were initiated, also without benefit (Figure 1A). Subsequent CSF pleocytosis with elevated B cells led to treatment with rituximab, but no improvement was observed. He continued to manifest worsening neurological symptoms, and repeat EMG/NCS test showed a diffuse neurogenic disorder with active denervation in two body segments, confirming both upper and lower motor neuron disease, clinically resembling ALS. Repeat CSF immunophenotyping showed elevated CD8 T and B cells (Figure 1D), indicating neurodegeneration with CNS immune dysregulation.

The patient’s clinical profile, disease progression, negative autoimmune workup, and lack of response to immunotherapies suggested that this novel *PIK3R1* mutation might have a direct role in his neurodegeneration.

### *PIK3R1* c.1710dup mutation causes haploinsufficiency

The *PIK3R1* c.1710dup mutation, located in exon 13, leads to frameshift and premature termination of all three isoforms of the encoded protein (Figure 2A). iPSC lines were generated from healthy donor and patient PBMCs. The *PIK3R1* mutation was corrected in the patient’s iPSCs using CRISPR-Cas9 as described. In addition, the *PIK3R1* c.1710dup was introduced into healthy control donor iPSCs using CRISPR-Cas9. Sanger sequencing of patient iPSCs confirmed the mutation and successful correction in the edited iPSCs (Figure 2B). Reverse transcription PCR (RT-PCR) directed at the mutation site showed less *PIK3R1* mRNA in patient iPSCs compared to corrected patient cells or healthy controls (Figure 2D). To determine whether reduced *PIK3R1* mRNA in patient iPSCs was due to mRNA decay we performed Sanger sequencing of *PIK3R1* cDNA from patient, corrected patient cells and healthy iPSCs. Only wild-type *PIK3R1* cDNA sequence was detected in patient-derived iPSCs, indicating complete mRNA decay of the mutated transcript (Figure 2E). Sanger sequencing of cDNA from healthy control iPSC into which we introduced *PIK3R1* c.1710dup also showed partial mRNA decay (Figure 2D). To examine the effect of mRNA decay on p85α protein levels, we immunoblotted iPSC lysates from our patient, corrected patient cells and healthy control cells. Levels of p85α protein were lower in patient’s iPSC lysates compared to those from the corrected patient cells or healthy control (Figure 2C). Following confirmation of mRNA decay in iPSCs, we differentiated iPSC into motor neurons (iMNs) using a three-step differentiation protocol [7, 8]. The terminally differentiated patient iPSC-derived motor neurons had normal morphology and expressed the motor neuron marker choline acetyltransferase (CHAT), but not the cortical neuron marker CTIP (Figure 2F). Sanger sequencing of cDNA extracted from terminally differentiated iMNs confirmed complete *PIK3R1* mRNA decay in the patient and engineered mutant iMN cells compared to corrected patient cells and healthy control cells (Figure 2G, H). These data confirm that the novel frameshift *PIK3R1* c.1710dup leads to p85α haploinsufficiency.

**Figure 2.**
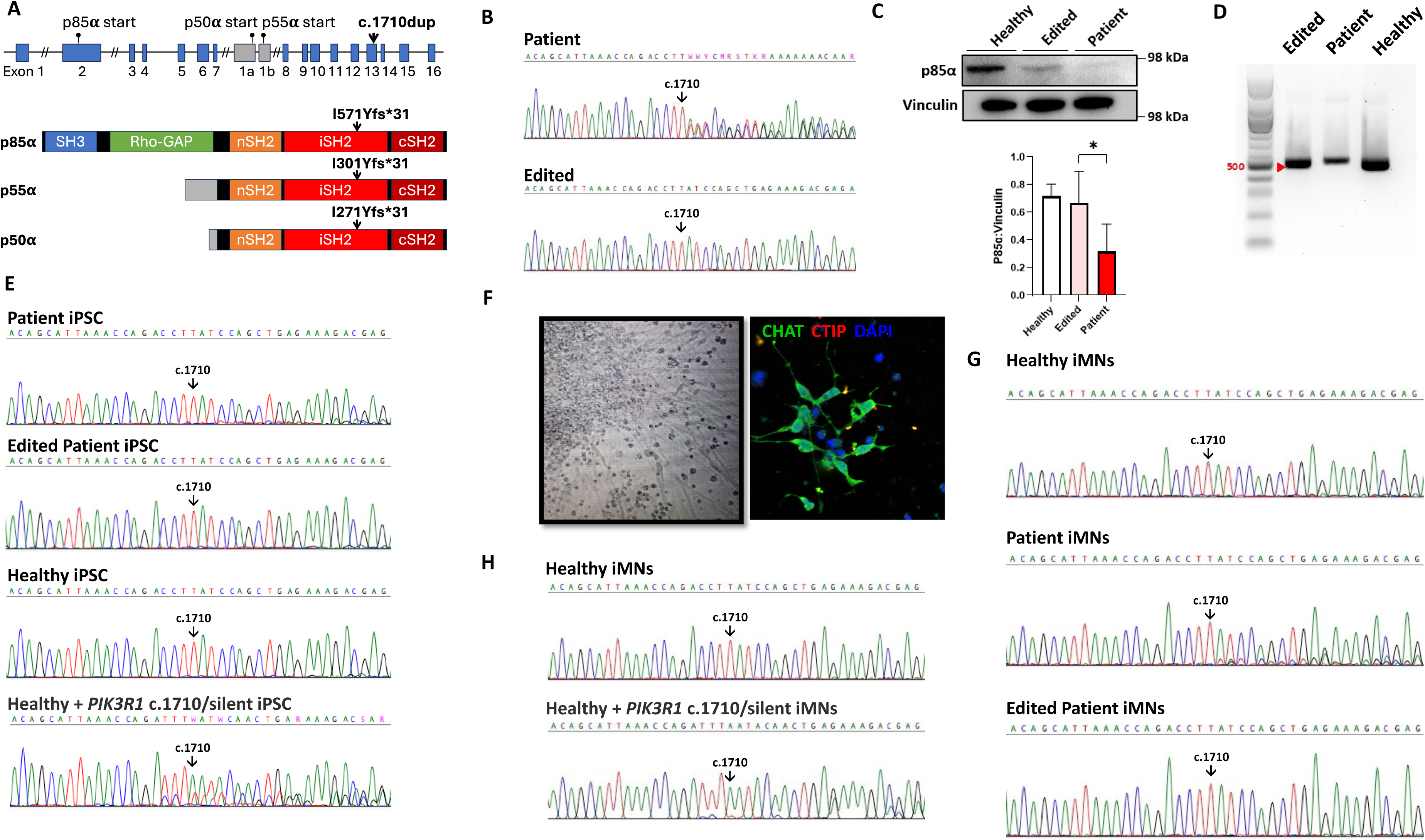
The effect of *PIK3R1* c.1710dup on *PIK3R1* mRNA and p85α protein. **A)** *PIK3R1* gene schematic representation showing the location of the *c.1710dup* mutation in exon 13 of *PIK3R1* and its predicted effects on the expression of *PIK3R1* encoded protein isoforms. **B)** Sanger sequencing of cDNA from patient iPSCs and base edited iPSCs showing wild-type *PIK3R1* sequence in the edited cell line, indicating correction of the mutation in patient’s iPSCs **C)** Immunoblot analysis revealed reduced p85α protein levels in patient-derived iPSCs compared to gene-corrected patient and healthy controls (Densitometry data are pooled from three independent experiments* p<0.05). **D)** RT-PCR analysis of mRNA using *PIK3R1* primers from patient, edited and healthy control iPSCs showing reduced expression of *PIK3R1* mRNA in patient iPSCs compared to edited and healthy control. **E)** Sanger sequencing of cDNA from patient iPSCs showing presence of wild-type *PIK3R1* sequence only, indicating complete degradation of the mutated transcript. **F)** Immunohistochemistry staining of terminally differentiated iPSC-derived motor neurons showed normal neuronal morphology in patient’s cells and expression of CHAT, a motor neuron marker, but not CTIP, a cortical neuron marker. **G)** Sanger sequencing of cDNA from terminally differentiated iMNs confirmed complete mRNA decay of *PIK3R1* c.1710dup transcript in patient’s compared to edited and healthy controls. **H)** Sanger sequencing of cDNA from terminally differentiated iMNs from healthy and *PIK3R1* c.1710dup introduced mutation showed only the normal mRNA sequence in both lines confirming an mRNA decay of the c.1710dup transcript.

### *PIK3R1* c.1710dup mutation activates the AKT pathway

Using iPSC and iMNs lysates from patient, edited patient, and control cells, we found the phospho-AKT (Thr308 and Ser473) to total AKT ratio was significantly elevated in iPSCs from the patient line compared to edited and control cells (Figure 3A). Measuring the pAKT (Thr308 & Ser473)/AKT ratio in iMN cell lysates showed the same trend in patient derived iMNs as in the healthy control (Supplemental Figure 2A). In order to assess AKT downstream signaling, we measured phosphoS6 and total S6 values and found no difference in phospho-S6/S6 ratios in iPSCs from patient, edited and healthy control (Figure 3B). The same measurement in iMN from patient and healthy donor cells demonstrated a decreased trend of pS6/total S6 ratio in patient cells (Supplemental Figure 2B). These data indicate that *PIK3R1* c.1710dup led to elevated phosphorylation on both pAKT (Ser473) and pAKT (Thr308) sites of AKT, and activated the AKT pathway with no effect on the pS6/S6 ratio.

**Figure 3.**
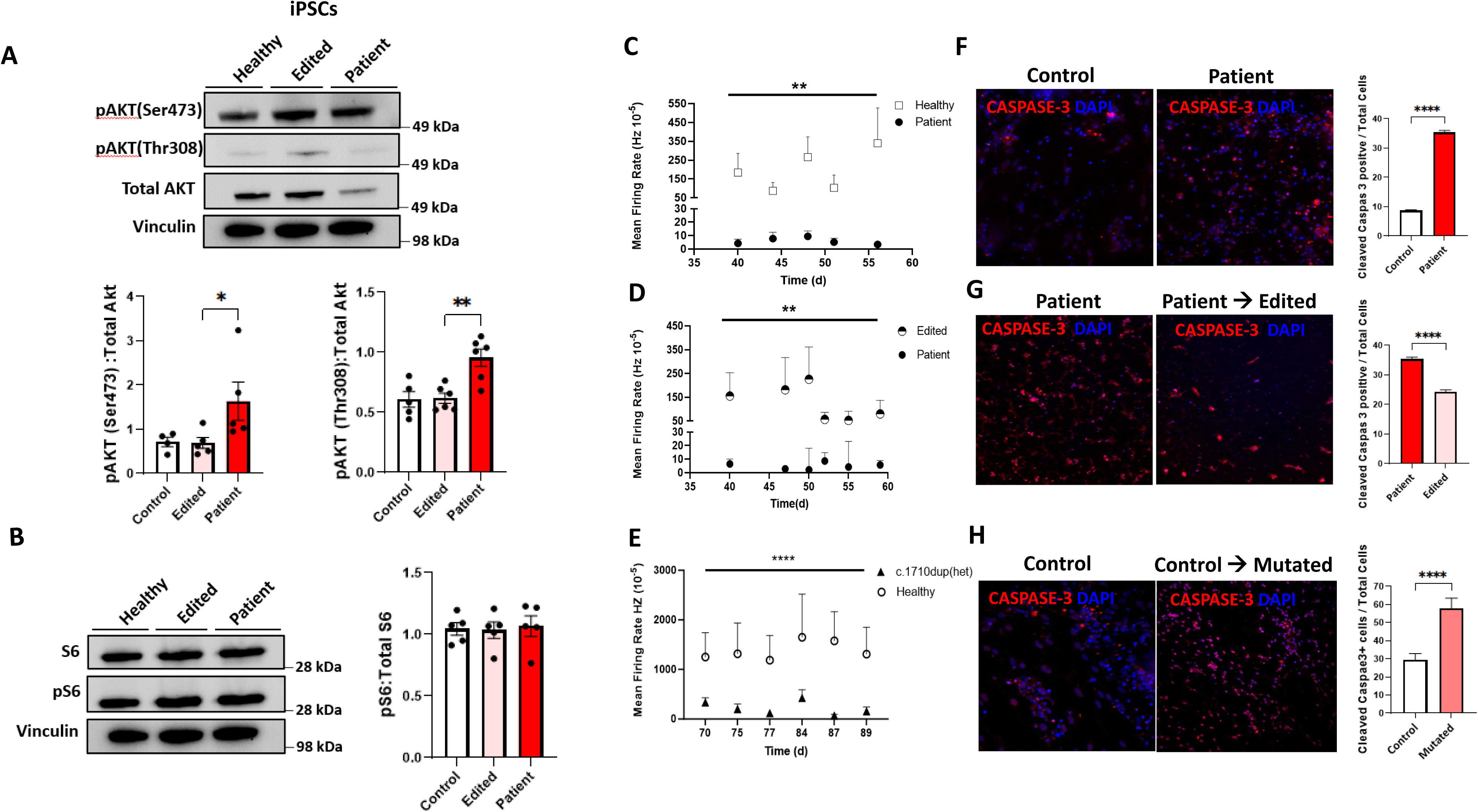
The effect of *PIK3R1* c.1710dup mutation on *PIK3R1* signaling pathway and neuronal electrical function and survival; **A)** Immunoblot analysis of pAKT (Thr308), pAKT (Ser473), total AKT from patient’s iPSCs lysate compared to edited and healthy controls are shown. Densitometry data are pooled from 5 (patient and healthy control) and 6 (patient and edited) independent experiments showed pAKT (Thr308) and pAKT(Ser473)/Total AKT ratio are significantly elevated in patient compared to edited and healthy controls ** p<0.01,* p<0.05 respectively. **B)** Pooled densitometry data from 4 independent immunoblot experiments of phospho-S6 to total S6 ratio did not demonstrate any change in patient derived iPSCs compared to healthy and edited cells. **C)** Multi-well microelectrode array (MEA) recordings of terminally differentiated motor neurons (iMNs) shows significantly reduced mean firing rate of patient-derived iMNs compared to healthy iMNs from day 32 to the end of the experiments (Data are representative of three separate experiments; ** p<0.01). **D)** MEA recordings shows normalized mean firing rate of iMNs from patient’s edited iPSCs compared to patient’s iMNs from day 32 to the end of the experiments (Data are representative of three separate experiments; ** p<0.01). **E)** CRISPR-Cas9 *PIK3R1* c.1710dup mutated iMNs exhibit reduced electrical activity compared to isogenic healthy control iMNs. (Data are representative of three separate experiments; **** p<0.0001). **F-G)** Immunohistochemistry staining for cleaved caspase-3 (Red) and DAPI (blue) in iMNs on Day 55 post differentiation show percentage of apoptotic cells was significantly increased in patient and *PIK3R1* c.1710dup mutated iMNs compared to healthy controls (Data are representative of three separate experiments ****p < 0.0001) and Gene correction significantly decreased cleaved caspase-3 expression (Red) to total number of cells in edited iMNs compared to patient cells. (Data are representative of three separate experiments ****p<0.0001). **H)** CRISPR-Cas9 *PIK3R1* c.1710dup mutated iMNs also showed significantly elevated cleaved caspase-3 positive cells compared to isogenic healthy control (Data are representative of three separate experiments ***p<0.001).

### Novel *PIK3R1* mutation leads to abnormal neuronal function and survival

To further investigate the role of *PIK3R1*c.1710dup effect in motor neurons, we differentiated iPSCs into iMNs and measured iMN spontaneous electrical activity and mean firing rate using electrode-embedded culture plates (multi-well microelectrode array (MEA) plates), starting on Day 30 post-differentiation (Pd) through Day 90 Pd. Patient iMNs had a significantly lower mean firing rate than healthy control iMNs (Figure 3C). Gene-corrected patient iMNs had normalized electrical activity compared to uncorrected patient iMNs (Figure 3D). To validate the effect of *PIK3R1* c.1710dup on motor neuron function and apoptosis, we differentiated iPSCs with engineered *PIK3R1* c.1710dup and their wild-type parental counterpart into iMNs. While both iPSC lines differentiated normally into motor neurons and showed normal morphology and expression of the motor neuron marker, CHAT, we found significantly reduced electrical activity in mutated iMNs compared to the wild-type parental cells (Figure 3E).

Abnormal electrical function in iMNs with *PIK3R1* c.1710dup led us to evaluate apoptosis. We examined fixed iMNs for Cleaved-Caspase3 and CHAT. Cleaved-Caspase3 expression was significantly increased in patient-derived iMNs on Day 56 Pd compared to the corrected patient cells and healthy control iMNs (Figure 3F, G).

Furthermore, caspase-3 expression in mutated iMNs was significantly elevated compared to control cells (Figure 3H), consistent with our findings in patient-derived iMNs. *PIK3R1* c.1710dup in terminally differentiated iMNs induced excess apoptosis implicating *PIK3R1* haploinsufficiency in neuronal function and survival.

### *PIK3R1* c.1710dup increases apoptosis in neuronal and Glial-like clusters of iMNs

Patient, healthy control, CRISPR-Cas9 engineered mutant and healthy parental strain iPSC-derived iMNs on Day 55 Pd were used for single cell RNA sequencing (scRNA-seq). Three clusters were identified based on cell marker genes for neuronal, glial-like, and cycling cells (described in methods and Supplemental Figure 2). Patient cells showed an increased percentage of neuronal, and a reduced percentage of glial-like cells compared to controls (Figure 4A, Supplemental Figure 2C). Pathway analysis in the neuronal cluster showed higher expression of stem cell differentiation, neural precursor differentiation, neuronal differentiation regulation, and axonogenesis in patient compared to healthy cells (Figure 4B). The intrinsic apoptosis signaling pathway and its regulation were also among the upregulated pathways in patient cells. These findings imply that neurogenesis and apoptosis pathways are highly activated in patient iMNs, indicating that the *PIK3R1* c.1710dup may drive increased neuronal differentiation and apoptosis in motor neurons (Figure 4B). Glial-like cells in the patient iMN culture also exhibited higher expression of pathways related to intrinsic apoptosis and their regulation, suggesting that *PIK3R1* c.1710dup may contribute to glial-like cell dysfunction and death, which may play roles in neurodegeneration (Figure 4C).

**Figure 4.**
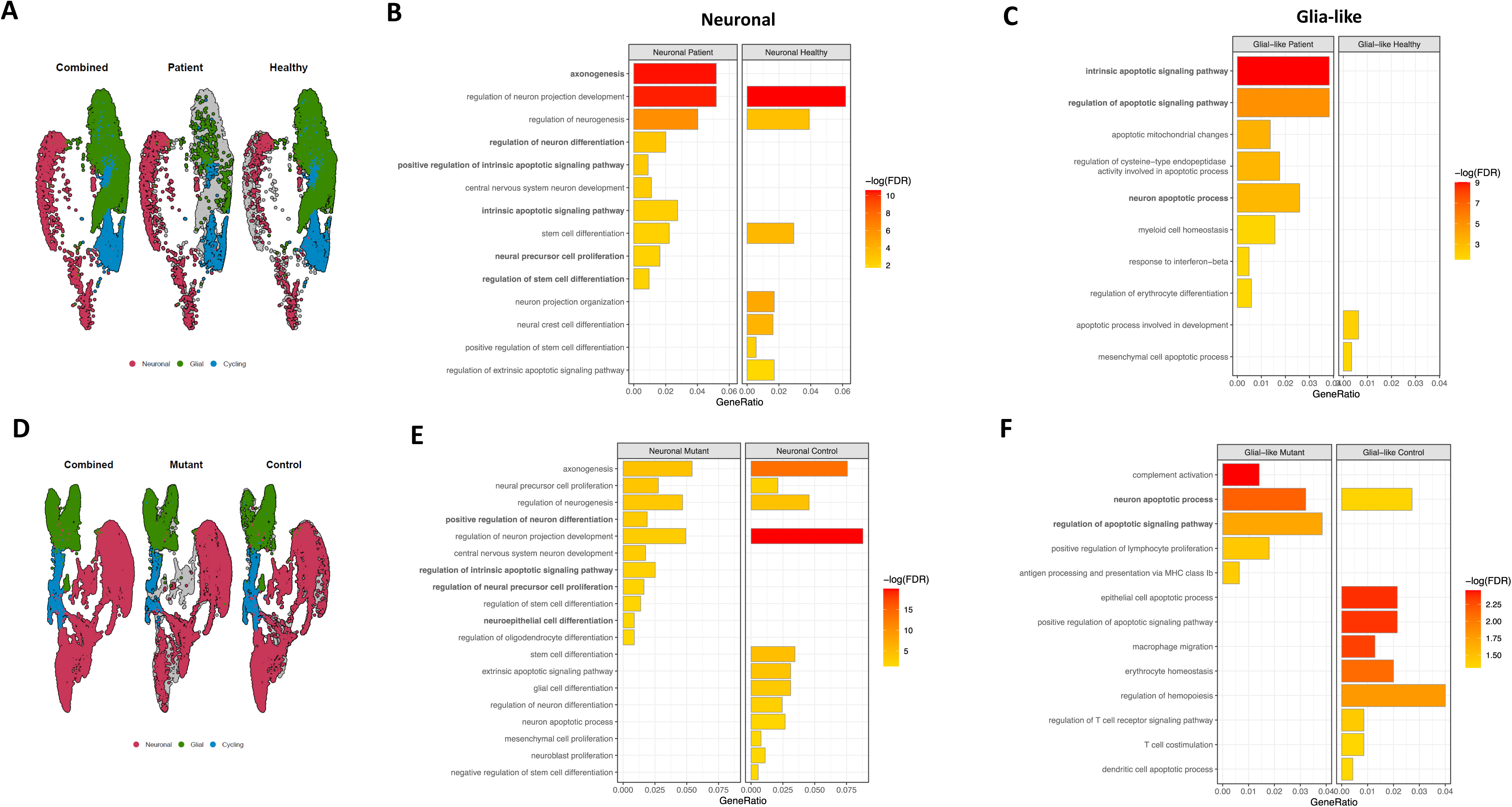
scRNA-seq analysis from patient and introduced *PIK3R1* c.1710dup mutation iMNs compared to control. **A)** UMAP plots shows three main clusters of cycling, neuronal and glia-like cells from patient and control iMNs. **B)** Differential gene expression analysis demonstrates upregulated apoptosis signaling pathways in neuronal clusters of patient iMNs compared to controls. **C)** Pathway analysis indicates upregulated apoptosis pathways in glia-like cluster from patient-derived iMNs compared to control. **D)** UMAP plots highlights neuronal, glia-like and cycling cells from CRSPR-CAS9 introduced *PIK3R1* c.1710 dup mutated and isogenic control iMNs. **E)** Differential gene expression and pathway analysis reveals upregulated intrinsic apoptosis signaling pathways in neuronal cluster of mutated iMNs compared to isogenic control. **F)** Pathway enrichment analysis indicating increased apoptosis-related pathways in glial-like cluster from mutant iMNs

We performed scRNA-seq on healthy control iMNs and on the same line with *PIK3R1* c.1710dup introduced. Three clusters of neuronal, cycling and glial-like cells were seen, just as above (Figure 4D). Pathway analysis in the neuronal cluster demonstrated a similar pattern of increased neurogenesis and intrinsic apoptosis pathways in the neuronal cell cluster from iPSCs with *PIK3R1* mutation (Figure 4E). Further evaluation of the glial-like cell cluster also showed higher expression of apoptosis signaling and its regulation pathways in mutant cells (Figure 4F), validating the effect of *PIK3R1* c.1710dup.

### *PIK3R1* c.1710dup reduces brain organoid size and accelerates apoptosis

Patient-derived, gene-edited, and healthy control iPSCs were differentiated into brain organoids as described. Organoids were imaged daily from Day 1 to Day 21 post-differentiation, reaching a plateau in size (Figure 5A). The size of each brain organoid was measured, and the mean organoid size was calculated for each group. Patient-derived organoids were significantly smaller than either gene-corrected or healthy control derived organoids, while no significant difference was observed between gene-corrected and healthy control derived organoids (Figure 5B). scRNA-seq was performed on brain organoids at six weeks of derivation from both patient-derived and gene-corrected lines. Cell cluster analysis based on gene expression identified cycling, neuronal, and glial-like populations, as described (Figure 5C). The neuronal cluster was more populous in patient-derived organoids, whereas glial-like cells were more abundant in gene-corrected organoids (Figure 5D). Gene expression analysis of each cell cluster in patient-derived organoids demonstrated increased expression of apoptosis-related genes compared to their gene-corrected counterparts (Supplemental Figure 3). Pathway analysis of enriched genes revealed that the intrinsic apoptosis signaling pathway was among the upregulated pathways expressed in both neuronal and glial-like clusters in patient organoids compared to gene-corrected organoids.

**Figure 5.**
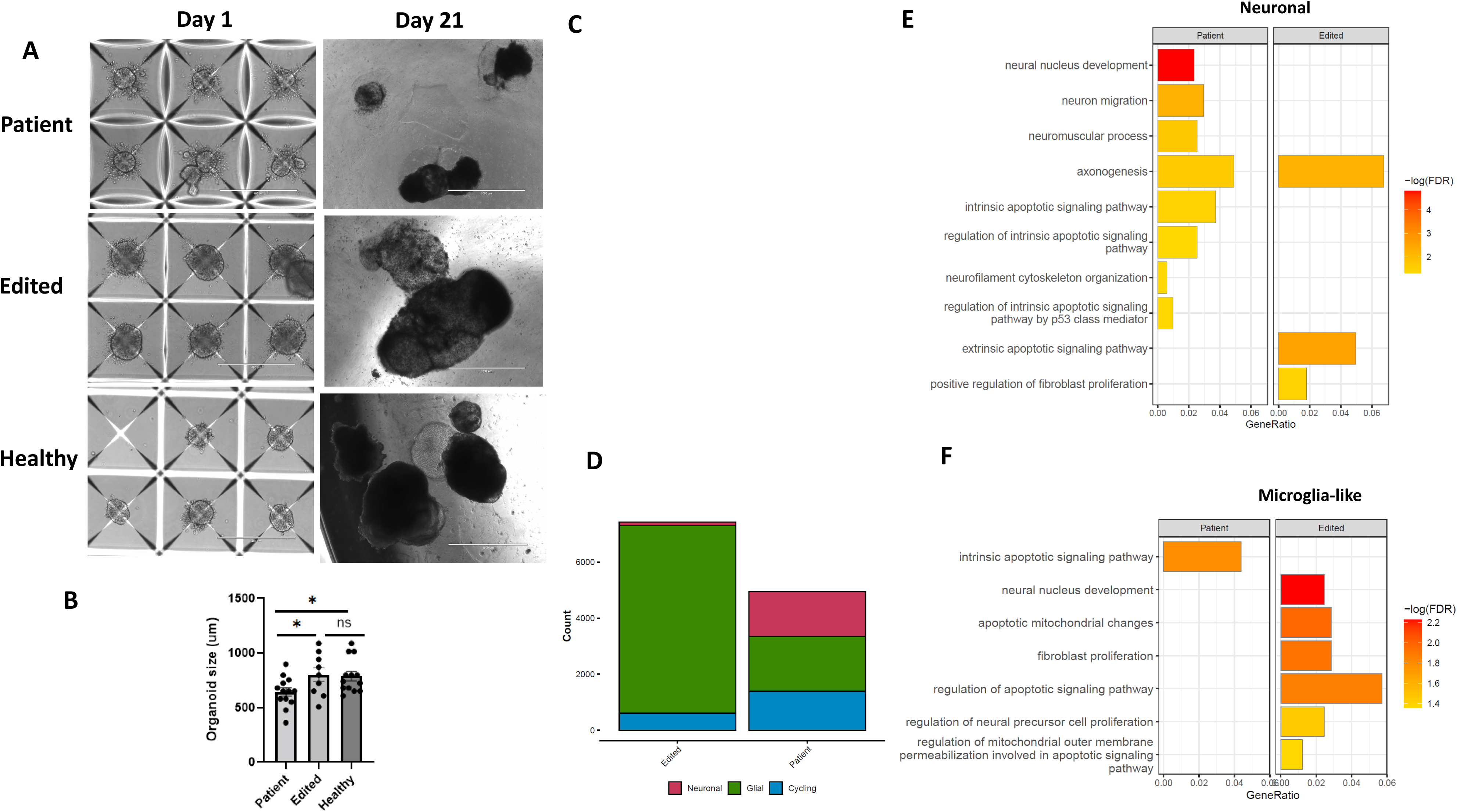
Patient and patient-edited iPSCs derived brain organoid differentiation and transcriptomics. **A)** Representative imaging of patient brain organoids shows smaller size in patient organoids compared to edited and healthy brain organoid. **B)** Measurement of mean size of brain organoids in patient showed significantly smaller size compared to healthy and edited organoids on Day 21. **C)** Analysis of data from single-cell sequencing of brain organoids shows neuronal, glia-like and cycling cells in patient and edited brain organoids. **D)** Graph-bar of each cluster in patient and edited organoids demonstrate larger neuronal and smaller glia cluster in patient derived organoids. **E)** Differential gene expression and pathway analysis reveals upregulated neurogenesis-associated and apoptosis signaling pathways in neuronal cluster of brain organoids from patient compared to control. **F)** Pathway enrichment analysis indicated increased apoptosis-related pathways in glial-like cluster from patient organoids.

## Discussion

We identified a patient with immunodeficiency and progressive motor neuron disease who carries a novel frameshift c.1710dup mutation in *PIK3R1*, resulting in *PIK3R1* haploinsufficiency. We provide the first direct evidence that *PIK3R1* haploinsufficiency impairs neuronal and glial function, ultimately contributing to neurodegeneration. This mechanism differs from previously reported *PIK3R1* mutations in APDS2, in which the mutant protein activates the p110δ catalytic subunit, leading to immune dysregulation, whereas in this case there is reduced protein expression [4, 9].

The patient’s early neurological presentation, combined with his immune deficiency and dysregulation, along with oligoclonal bands (OCBs) and an elevated IgG index, initially led to treatment with immunomodulatory therapies. However, his evolving clinical features led to a diagnosis of slowly progressive ALS. CSF immunophenotyping showed an elevated percentage of CD8 T cells and B cells, while myeloid cells were reduced compared to healthy controls (Figure 1D). These findings suggested a potential link between systemic immune dysregulation, CNS immune activation, and neurodegeneration. OCBs and elevated IgG indices have been observed in a subset of ALS patients, particularly those with genetic ALS compared to sporadic cases. OCBs have been attributed to a localized immune response, intrathecal IgG synthesis by B cells within meningeal tertiary lymphoid structures, and to a potentially compromised blood-brain barrier [10]. More effector T cell profiles in the blood and CSF of ALS patients may correlate with poorer prognosis [11]. However, the significance of these findings in motor neuron disease progression and its underlying mechanisms remain unclear [12]. Immunological findings like OCBs and an elevated CSF IgG index may help clinicians diagnose immune mediated neurological disorders, but they do not differentiate neuroinflammatory from neurodegenerative disorders.

We investigated the underlying mechanism by which *PIK3R1* c.1710dup causes neurodegeneration. PIK3R1 haploinsufficiency enhanced AKT phosphorylation in patient-derived iPSCs and iMNs (Figure 3A, C), while pS6 remained unaltered (Figure 3B,D), indicating distinct downstream signaling alterations in these cells. Heterozygous *PIK3R1* knockout mice with reduced expression of p85α and its alternative isoforms exhibited increased AKT phosphorylation and activity in hepatocytes [13]. Moreover, PIK3R1 loss has been shown to induce aberrant AKT activation via both p110-dependent and independent mechanisms [14, 15]. However, these effects are tissue-specific, as PIK3R1 deficiency in myocytes results in decreased AKT activation [16].

The PI3K/AKT pathway is a critical regulator of neuronal development, synaptic plasticity, axonal repair, myelination, and apoptosis [17]. AKT signaling plays a central role in promoting cell proliferation and survival by inhibiting pro-apoptotic molecules and it is a key regulator of neuronal proliferation [18]. However, AKT activation is a double-edged sword, increasing susceptibility to oxidative stress and cell death under certain cellular conditions [19]. Both patient-derived and CRISPR-engineered *PIK3R1*-mutant iMNs and brain organoids exhibited increased neuronal precursor proliferation and neurogenesis, suggesting that *PIK3R1* haploinsufficiency may drive early-stage differentiation through AKT activation. However, elevated caspase-3 expression in terminally differentiated iMNs, along with scRNA-seq demonstrating heightened apoptosis-related pathways in neuronal and glia-like populations in both patient-derived and mutant iMNs and brain organoids, suggest that *PIK3R1* haploinsufficiency may also activate proapoptotic signaling pathways. Excessive AKT activation enhances oxidative stress and increases susceptibility to reactive oxygen species (ROS)-mediated cell death, which may contribute to our findings [20]. Additionally, the PI3K/AKT pathway regulates multiple cellular functions, and dysregulation caused by PIK3R1 haploinsufficiency may modulate FOXO transcription factors, reduce antioxidant protein levels, and promote proapoptotic signaling, leading to oxidative stress-induced cell death [21, 22]. PIK3R1 can exert AKT-independent functions in cells and p85α knockout or knockdown cells exhibit increased nuclear accumulation of X-box binding protein-1 (XBP-1) and elevated apoptosis [23].

Electrical activity of iMNs derived from patient and mutant cells was significantly lower than that in cells derived from healthy controls. Further, gene correction restored normal activity, confirming that *PIK3R1* haploinsufficiency disrupts neuronal function. Abnormalities in neuronal electrical activity can increase apoptosis rates, which may explain our findings [24, 25].

Taken together, our findings provide a novel aspect of *PIK3R1* function and underscore its essential role in neuronal integrity and survival. Further research will elucidate the precise molecular mechanisms by which *PIK3R1* haploinsufficiency may impair neuronal and glial function and survival, providing critical new insights into neurodegenerative diseases.

## Methods

### Sex as a biological variable

For studies involving humans and human samples, sex was not considered as a biological variable.

#### CSF flow cytometry

CSF from patient and healthy donors were immediately transferred to lab following lumbar puncture and spun down in 300GX for 10 minutes. Cell free CSF was collected, and cell pellets were used for flow cytometry. Cells were stained for 35 surface markers as described in Supplemental Table 1 and run with Cytek Machine Cytek Aurora 5 Laser UV/V/B/YG/R (64 + 3 Channel). Data was analyzed through Cytobank software.

### Reprogramming and maintenance of iPSCs

Healthy control iPSCs were generated from healthy volunteer peripheral blood CD34^+^ hematopoietic stem/progenitor cells using previously described methods [26]. Patient and healthy donors iPSCs were generated and provided by the iPSC Core Facility of the National Heart, Lung, and Blood Institute (NHLBI) using methods detailed in a prior publication [27]. The human iPSCs were cultured in E8 Flex medium (Thermo Fisher Scientific) or mTeSR-Plus medium (STEMCELL Technologies) in a 5% CO_2_ incubator at 37 °C on plates coated with ESC-qualified Matrigel (Corning).

### iPSC derived motor neuron differentiation

Motor neurons were differentiated from iPSCs following published literature [28]. Briefly, low density of iPSCs colonies were seeded on a Matrigel-coated 6 well plate in E8 Flex medium which was then replaced by Stage 1 iMN media for 6 days. 2 ml per well of fresh media were changed every other day. On day 6, the resulting neuroepithelial cells were dissociated with accutase treatment (Thermo Fisher) and collected by centrifugation at 1000 rpm for 5 min and re-plated at a density of ∼6e5 cells/well in a 6-well plate coated with Matrigel in Stage 2 media. The cells were continuously cultured in Stage 2 media for 6 days with media change at every other day. After that the media was replaced with Stage 3 media for neuronal maturation with media change every other day until 20 days post-differentiation. Stage 1media contains Iscove Modified Dulbecco Media (IMDM) (Gibco) and Ham’s F12 (Gibco) media mixed in a 1:1 ratio with 1% Non-Essential Amino Acids (NEAA) (Gibco), 1X B27 supplement (Gibco), 1X N2 supplement (Gibco), 1% antibiotic-antimycotic (Gibco), 0.2 µM LDN193189 (Cayman Chemical), 3 µM CHIR99021 (Xcess bioscience), and 10 µM SB431542 (Cayman Chemical). Stage 2 media were made with Stage 1 media plus 0.1 µM All-trans retinoic acid (Stemgent) and 1 µM Sonic Hedgehog Agonist SAG (Cayman Chemical).. Stage 3 media contained 1:1 IMDM (Gibco) and F12 media (Gibco), and 1% NEAA (Gibco), 1X B27 (Gibco), 1X N2 (Gibco), 1% antibiotic-antimycotic (Gibco), 0.1 µM SAG (Cayman Chemical), 0.5 µM all-trans retinoic acid (Stemgent), 0.1 µM compound E (Calbiochem), 2.5 µM DAPT (Cayman Chemical), 200 ng/ml Ascorbic Acid (Sigma), 10 ng/ml BDNF (PeproTech), 10 ng/ml GDNF (PeproTech). Cells were maintained in Stage 3 media with media change every other day until 20 days post-differentiation.

### Brain organoid differentiation and preparation

Brain organoids containing endothelial cells were derived from human iPSCs based on our established protocol with some modification [29].Briefly, human iPSCs were cultured on Matrigel in E8 Flex medium in 6 well plates until reaching ∼80% confluence. The iPSCs were dissociated into single cells to make embryoid bodies (EB) using Aggrewell 400 plate (STEMCELL Technologies) following the manufacturer’s instruction. 24 h later, 10-20 formed EBs were transferred per well into the low attachment 24 well plate in 1 ml of E8 flex medium. 72 h later, 500 µl of medium was removed and the EBs were treated with 15 cold Matrigel by adding directly onto the medium. After 4 h of incubation, 500 µl of DMEM/F12 supplemented with B27 and Activin A(125ng/ml) was added on top of the cells. 24 h later, half of the media were replaced with E8 Flex containing 10 ng/ml BMP4 and Matrigel overlay was performed as above again. After 72 h, half of the medium was replaced with DMEM/F12 containing B27, 1 mM 8bromo-cAMP, 100ng/ml VEGF and antibiotics. Half of the medium were changed with the cerebral organoid maturation medium (STEMCELL Technologies) 72 h later and every other day till the end of experiments.

### CRISPR-Cas9 editing of iPSCs

For correction of the PIK3R1 c.1710dup mutation, patient iPSCs were electroporated with 1 µg of Cas9 nuclease mRNA and 10 µg (∼60 pmol) of chemically-modified synthetic guide RNA (U*U*C*UCAGCUGGAUAAAGGUC + Synthego modified EZ Scaffold; Synthego) as well as 2.5 µg of a symmetrical single-stranded oligodeoxynucleotide (ssODN) donor DNA (Integrated DNA Technologies; IDT) for correction of the patient mutation (ssODN sequence: 5’-C*T*GAGTATCGAGAAATTGACAAACGTATGAACAGCATTAAACCAGACCTTATCCA GCTGAGAAAGACGAGAGACCAATACTTGATGTAAGTATTTGA*A*A-3’; asterisks denote phosphorothioate bond modifications for enhanced oligo stability) by homology-directed repair at the CRISPR-Cas9 target site. 11.4 µg of i53 mRNA (48.6 pmol; CELLSCRIPT) was also included in the electroporation mixture to enhance homology directed repair by inhibiting 53BP1 in the non-homologous end-joining pathway [30].Cell were electroporated in a total volume of 50 µL of MaxCyte electroporation buffer (MaxCyte) in a MaxCyte ATx electroporation system with program iPSC 3. Following electroporation, iPSC clones were expanded and genomic DNA was screened by Sanger sequencing to identify clones with correction of the heterozygous c.1710dup mutation.

For targeted heterozygous knock-in of the PIK3R1 c.1710dup mutation, healthy donor iPSCs were electroporated with Cas9 ribonucleoproteins consisting 100 pmol of S.p. HiFi Cas9 nuclease V9 protein (IDT) and 500 pmol of chemically-modified synthetic guide RNA (U*U*U*CUCAGCUGGAUAAGGUC + Synthego modified EZ Scaffold; Synthego) as well as 10 µg each of two asymmetrical ssODN donor DNAs (IDT) for homology-directed repair at the CRISPR-Cas9 target site. One ssODN contained the PIK3R1 c.1710dup mutation for knock-in (5’-A*G*ACTTGAAGAAGCAGGCAGCTGAGTATCGAGAAATTGACAAACGTATGAACAG CATTAAACCAGACCTTTATCCAGCTGAGAAAGACGAGAGACCAATACT*T*G-3’; underlined nucleotide corresponds to c.1710dup; asterisks denote phosphorothioate bond modifications for enhanced oligo stability) and the other ssODN contained silent mutations to the guide RNA target site (A*G*AAGACTTGAAGAAGCAGGCAGCTGAGTATCGAGAAATTGACAAACGTATGAA CAGCATTAAACCAGATTTAATACAACTGAGAAAGACGAGAGACCAATACTTGA*T*G; underlined nucleotides denote silent mutations; asterisks denote phosphorothioate bond modifications) to enable insertion of the c.1710dup mutation at one allele while protecting a healthy PIK3R1 sequence from Cas9-induced indels at the other allele. 10 µg of i53 mRNA (42.6 pmol) was also included in the electroporation mixture to enhance homology directed repair. Cells were electroporated in 100 µL of homemade buffer (Opti-MEM + 7.25 mM ATP + 11.8 mM MgCl2) using an Amaxa 4D Nucleofector system (Lonza) with program CA-137. Following electroporation, iPSC clones were expanded and genomic DNA was screened by Sanger sequencing to identify clones with simultaneous heterozygous knock-in of the c.1710dup mutation at one allele and the protective silent mutations at the other allele.

### RT-PCR and sequencing of PIK3R1 cDNA

Whole RNA was purified from iPSCs or iMNs using RNeasy Plus Mini kit (QIAGEN), and cDNA was synthesized from 100-500 ng of RNA using SuperScript III First-Strand Synthesis SuperMix (Thermo Fisher Scientific) with a 1:1 mixture of oligo(dT) and random hexamers. Half of the cDNA synthesis reaction was used as template for PCR with P1185_f (5’-CGCACATCCCAGGAAATCCAAATG-3’) and P1186_r (5’-GCAACAGGTTTTCAGCTTTGTTTCG-3’) primers specific for PIK3R1 cDNA. Purified PCR products (NucleoSpin Gel & PCR Clean-up kit; Macherey-Nagel) were used as templates for Sanger sequencing with P1185_f primer to assess the presence of healthy versus c.1710dup mutation PIK3R1 transcripts.

### Electrophysiological Analysis Using Microelectrode Array

Human neurons (200,000 cells/well) were seeded into a 48-well microelectrode array (MEA) plates (M768-tMEA-48B; Axion BioSystems) containing 16 active recording electrodes/well. Electrophysiological activity, noted by increased spike rate in the wells, increased significantly by 21 days in vitro and was monitored by recording spontaneous electrical activity in all wells for 5 minutes per day to establish a baseline. Spontaneous electrical activity was recorded 3-4 days per week. Quantitation of electrical activity (number of spikes, mean firing rate, and number of bursts) was completed with Axion BioSystems AxIS Integrated Studio 2.5.2 software.

### Immunohistochemistry Staining and image analysis

Motor neurons were fixed with 4% paraformaldehyde diluted in PBS for 10 minutes. Motor neurons were blocked with 10% normal donkey serum, 0.5% Triton-100, 98% (Cat # 327371000) diluted in PBS for 1 hour at room temperature. Cells were incubated in primary antibodies (prepared in BOND Primary Antibody Diluent (REF #AR9352)) overnight at 4 degree C (1:400 Cleaved Caspase-3 (D175) Rabbit Ab (REF # 9661S); 1:500 Anti-Cholin Acetyltransferase (REF # AB144).

Motor neurons were washed with PBS three times, followed by a 1-hour incubation in secondary antibody diluted in BOND Primary Antibody Diluent (1:500 Alexa Fluor 594 anti-rabbit (REF# A11012); 1:500 FITC anti-Goat (REF # A16000) and then washed three times with PBS. Motor neurons were then incubated in DAPI (1:1000) diluted in BOND Primary Antibody Diluent for 10 minutes. Following three washes with PBS, images were acquired as described below.

Neuronal cultures (20,000 cells/well) were seeded into 96-well plates and maintained at 37°C with 5% CO2. Following treatment, the neurons were fixed and stained as described above with DAPI cell nuclei, Caspase 3 with red color and ChAT with green color.The cells were imaged with the Molecular Devices (Sunnyvale, CA) ImageXpress Micro Laser Confocal imager, acquiring 4 images/well at 10x magnification with a 50 µm slit spinning disk confocal mode. High content imaging/analysis was achieved with MetaXpress v. 6.7.2.290 analysis software for neuronal cell counts and neurite outgrowth. Neuronal viability and neurite length were quantitated and plotted with GraphPad Prism 7.02 (GraphPad Software).

### Western Blot

Western blot experiments was done on cell pellets as previously described [31]. Briefly media was removed and following a wash with 1 x PBS (REF #14190-144), replaced with fresh media. Using cell scraper (REF #83.3951), cells were collected in a sterile tube and spun at 600 g for 5 minutes, resuspend in 1ml media again and spun one more time in microcentrifuge tube at 600 g for 5 minutes. The supernatant was removed and cell pellets were put on dry ice (2-3 minutes) then stored at −80 degrees C. RIPA Buffer (with protease/phosphatase inhibitors) was added to the cell pellet. Cell pellets were incubated on ice for 20 min. Using centrifugation, the whole cell lysates were separated from cell debris at 10,000 x g at 4 C for 15 min. Pierce BCA Protein Assay Kit (Doc. Part No. 2161296 / Pub No. MAN0011430) was used to determine the protein concentration.

Laemmli buffer (Tris-Glycine SDS) was added to lysate samples. The samples were heated at 95 C for 3 minutes and centrifuged. Samples were loaded into a NuPAGE 4-12% Bis-Tris Gel (REF # NP0336BOX) with a PageRuler Plus Prestained Protein Ladder (Catalog Number: 26619) and separated at 160 V. The gels were transferred onto Immobilon Polyvinylidene fluoride (PVDF) membranes (REF # IB24002) at 20 volts for 7 minutes. Five percent milk in PBS containing 0.05% Tween-20 (PBS-T) was used to block the protein membranes for 30 minutes at room temperature. A primary antibody in BSA was added to the membranes, which were then incubated overnight at 4 C. Primary antibodies used included: vinculin (Abcam; Cat # AB129002), Akt Rabbit Ab (Cell Signaling Technology; Cat # 9272S), phospho-Akt (Ser473) Rabbit mAb (Cell Signaling Technology; Cat #4060S), phospho-Akt (Thr308) Rabbit mAb (Cell Signaling Technology; Cat #4056S), S6 ribosomal protein Rabbit mAb (Cell Signaling Technology; Cat #2217S), phospho-S6 ribosomal protein (Ser240/244) Rabbit mAb (Cell Signaling Technology; Cat #5364S), PI3 kinase p85 alpha Rabbit mAb (Cell Signaling Technology; Cat # 4257). Using PBS-T, the membranes were washed three times at 5 min per wash. A complementary HRP-conjugated secondary antibody was added and the membrane was incubated for 1 h at room temperature. The membranes were then washed three times with PBS-T at 10 min per wash. Super Signal West Femto Maximum Sensitivity Substrate was used to activate HRP luminescence. The Molecular Imager ChemiDoc Touch system (Bio-Rad) was used to develop the membranes.

### scRNA-seq library preparation and analysis of iMNs

Two batches of single cell RNA libraries were made using the 10X Genomic Single Cell Chromium 3’ mRNA kit version 3 and sequenced on either a NextSeq 500 (first batch) or NovaSeq 6000 sequencer (second batch). The first batch was the mutated sample and matching control; the second had the patient and healthy samples. All libraries had over 450 million reads. The Q30 values across the four libraries averaged 96.4%, 95.0%, and 96.3% for the barcode, RNA read, and UMI, respectively. Initial processing was accomplished using cellranger (10x Genomics), which included alignment to the human reference GRCh38-2020-A with the include introns parameter set to True. Mutated and control samples were processed with cellranger-7.1.0; patient and healthy with cellranger-8.0.0. Median gene count range was 64-114 thousand with sequencing saturation between 37-61% and 70-77% mapping to the transcriptome. The fraction of reads in cell was between 77-86%. Other key QC values are listed in the table (Supplemental Figure 2A)

Following Cell Ranger processing, the ‘isOutlier’ function from the R package scuttle v1.12.0 was used to identify cells that deviated by at least three median absolute deviations from the median number of reads, number of features, and percent mitochondrial reads. Seurat v4.4.0 was then used to filter outlier cells based on the ‘isOutlier’ results, along with additional stringent cutoffs to enhance consistency between samples. Specifically, cells were retained only if the mitochondrial percentage was below 15%, the number of features was below 13,000, and above 1,000. After filtering, data was log-normalized using the Normalize Data function. Following normalization, samples were CCA-integrated to their matching control (patient and healthy; mutant and control) using the FindIntegrationAnchors function.

To define cell types, the mutant and control integrated object was divided into seven clusters using nPC = 20 and a resolution of 0.1. Clusters were assigned labels as “cycling”, “glial”, and “neuronal” based upon key marker genes. SMC4, a subunit of the condensin complex, and CENPF, a centromeric protein, were used as markers to identify cycling cells. Glial (mesoderm-like) cells were identified by expression of TWIST1 and FOXC1. ELAVL3, CHAT, and STMN2 were used as markers for neuronal cells. A second integration between the patient/healthy and mutant/control integrated objected was used to transfer these labels from the mutant/control set to the patient/healthy sample set. UMAPs and dot plots were created with SCpubr v 2.0.2.(Supplemental Figure 2B)

Marker genes for neuronal and glial cells were identified using the ‘FindAllMarkers’ function with the MAST test and the following parameters: only.pos = TRUE, min.pct = 0.25, and logfc = 0.15. Genes with an adjusted p-value less than 0.1 were considered significant. Pathway analysis was performed using clusterProfiler v4.10.0 and GO.db v3.18.0. Related pathways were collapsed using the clusterProfiler simplify function and a cutoff of 0.7. Barplots were created with ggplot2 v 3.4.4. Pathways presented in the visualizations were selected from the full list of differentially expressed pathways using a custom script. Key search terms included, but were not limited to: “neuro*”, “apop*”, and “immun*”.

### Brain organoid scRNA-seq analysis

Single cell RNA libraries of edited and patient sample were made using 10X Genomic Single Cell Chromium 3’ mRNA kit version 3 and sequenced on NextSeq 500. Patient and edited libraries had over 77 and 97 million reads, respectively. The Q30 values for patient sample were 96.4%, 87.7%, and 94.9% for the barcode, RNA read, and UMI, respectively. The Q30 values for edited sample were 96.3%, 89.2%, and 95% for the barcode, RNA read, and UMI, respectively. Initial processing was accomplished using cellranger 7.0.0, which included alignment to the human reference GRCh38-2020-A with the include introns parameter set to True. Median genes per cell were 1,159 and 596 and sequencing saturation were 46.3% and 49.7% for patient and edited libraries, respectively.

The downstream analysis following Cell Ranger processing was performed in Seurat v4.1.1. Cells with number of features lower than 300 and higher than 4000 and mitochondrial reads higher than 20% were filtered out. Doublets detected by doubletFinder_v3 with pN = 0.25, pK = 0.24 and other default parameters were filtered out. After filtering, data was log-normalized using the NormalizeData function. Following normalization, samples were CCA-integrated using the FindIntegrationAnchors function.

Celltype annotation was performed using SingleR. The “Reproducible Brain Organoids” database from Human Cell Atlas of Broad Institute were used as reference. Cluster markers and DE genes were identified using FindMarker function in Seurat with the MAST test and the following parameters: only.pos = TRUE, min.pct = 0.25, and logfc = 0.15. Genes with an adjusted p-value less than 0.1 were considered significant for the purpose of pathway analysis. The pathway analyses were performed using clusterProfiler v4.12.0 and GO.db v3.19.1. Related pathways were collapsed using the clusterProfiler simplify function and a cutoff of 0.7. Barplots were created with ggplot2 v3.5.1. Pathways represented in the visualizations were selected from the full list of differentially expressed pathways using a custom script. Key search terms included but not limited to: “neuro*”, “apop*”, and “immun*”. Heatmaps were created with ComplexHeatmap v2.21.2.

### Statistics

*P* values and FDR *q* values resulting from 2-tailed tests were calculated using statistical tests stated in the figure legends. using GraphPad Prism v10.1.0. Differences between groups with *P* or *q* values < 0.05 were considered statistically significant

### Data Availability

All single-cell data included in this manuscript has been submitted to dbGAP under the accession XXXXX (to be provided upon acceptance). All code used for initial processing is available in the cell-seek pipeline repository (https://github.com/OpenOmics/cell-seek) and code for the downstream analysis is available on Github here: link to be provided.

### Study approval

Peripheral blood and CSF from patient and healthy donors were obtained after written informed consent following the Declaration of Helsinki. All human studies were approved by the NIAID and NINDS Institutional Review Board.

## Authors Contributions

BC and CLS share the first author position. BC and CLS contributed in conducting experiments, designing experiments, acquiring data, analyzing data, writing the manuscript. JS, TW, VM, LH, SD, SSDR, BP, conducted experiments, designing experiments, acquiring data. TEM and SP analyzed data and writing the manuscript. KY acquired and analyzed data and writing the manuscript. BG,CZ acquired clinical data. LDN, SMH and AN contributed in designing experiments, data interpretation and writing the manuscript. FS contributed in conducting experiments, designing experiments, analyzing and interpreting data and writing the manuscript

## Acknowledgment

We thank Mehrnoosh Abshari and Elina Stregevsky at combined technical research core in national institute of dental and craniofacial research (NIDCR) for their help in spectral flow cytometry. We are also so grateful to Alfred Killy, Francisco Otaizo-Carrasquero and Genomic Research Section Research Technologies Branch at NIAID for their essential help to perform scRNA-seq and Jizhong Zou and iPSC core team at NHLBI for their assistance in reprogramming of peripheral blood to iPSCs. This work utilized the computational resources of the NIH HPC Biowulf cluster. (http://hpc.nih.gov). This research is supported by Division of Intramural Research of NIAID, NIH and intramural research program of NINDS, NIH.

---

**Supplemental Figure 1. Peripheral blood immunophenotype and CSF findings in a patient with a *PIK3R1* c.1710dup mutation. A)** Peripheral blood immunophenotyping showing normal counts of CD4+ T cells, CD8+ T cells, and CD19+ B cells. **B)** CSF analysis demonstrates intermittent lymphocytic pleocytosis, elevated IgG levels, and isolated OCBs.

**Supplemental Figure 2. The effect of *PIK3R1* c.1710dup mutation on *AKT* signaling.**

Immunoblot analysis of pAKT (Thr308), pAKT (Ser473), total AKT from patient’s iMNs lysate compared to edited and healthy controls are shown. **A)** Immunoblots from iMNs in patient nd healthy are shown. Densitometry data from pooled three independent experiments confirmed elevated pAKT (Thr308 and Ser473)/Total AKT ratio in patient’s iMNs compared to healthy controls. **B)** Pooled densitometry data from 4 independent immunoblot experiments of phospho-S6 and total S6 showed patient derived iMNs have decrease pS6 to Total S6 ratio compared to healthy cells.

**Supplemental Figure 3. scRNA-seq analysis of iMNs with *PIK3R1* c.1710dup mutation**

**A)** Variable quality control metrics across scRNA-seq data from iMNs. **B)** Marker gene expression dot plots of all iMN scRNA-seq samples. **C)** Proportion bar plot showing percentage of cells in each cluster for the patient iMN and matching control.

**Supplemental Figure 4. scRNA-seq analysis of brain organoids with *PIK3R1* c.1710dup mutation. A)** Heatmaps show differentially expressed genes for neuronal and **B)** glial-like cell types. For visualization, genes with an adjusted p-value less than 0.1 were selected and the apoptotic pathway genes are annotated.

**Supplemental Table 1. Antibodies for immunophenotyping of CSF using spectral flow cytometry are shown.**

## References

1. Kriplani, N., et al., Class I PI 3-kinases: Function and evolution. Adv Biol Regul, 2015. 59: p. 53–64.

2. H, I.J., et al., Hyperactivation of the PI3K pathway in inborn errors of immunity: current understanding and therapeutic perspectives. Immunother Adv, 2024. 4(1): p. ltae009.

3. Innes, A.M. and D.A. Dyment, SHORT Syndrome, in GeneReviews(®), M.P. Adam, et al., Editors. 1993, University of Washington, Seattle Copyright © 1993-2025, University of Washington, Seattle. GeneReviews is a registered trademark of the University of Washington, Seattle. All rights reserved.: Seattle (WA).

4. Lucas, C.L., et al., Heterozygous splice mutation in PIK3R1 causes human immunodeficiency with lymphoproliferation due to dominant activation of PI3K. J Exp Med, 2014. 211(13): p. 2537–47.

5. Dyment, D.A., et al., Mutations in PIK3R1 cause SHORT syndrome. Am J Hum Genet, 2013. 93(1): p. 158–66.

6. Patel, V., W. Cui, and J.M. Cobben, SHORT syndrome with microcephaly and developmental delay. Am J Med Genet A, 2023. 191(3): p. 850–854.

7. Baxi, E.G., et al., Answer ALS, a large-scale resource for sporadic and familial ALS combining clinical and multi-omics data from induced pluripotent cell lines. Nat Neurosci, 2022. 25(2): p. 226–237.

8. Workman, M.J., et al., Large-scale differentiation of iPSC-derived motor neurons from ALS and control subjects. Neuron, 2023. 111(8): p. 1191–1204.e5.

9. Deau, M.C., et al., A human immunodeficiency caused by mutations in the PIK3R1 gene. J Clin Invest, 2014. 124(9): p. 3923–8.

10. Ticozzi, N., et al., Oligoclonal bands in the cerebrospinal fluid of amyotrophic lateral sclerosis patients with disease-associated mutations. J Neurol, 2013. 260(1): p. 85–92.

11. Yazdani, S., et al., T cell responses at diagnosis of amyotrophic lateral sclerosis predict disease progression. Nat Commun, 2022. 13(1): p. 6733.

12. Klose, V., et al., CSF oligoclonal IgG bands are not associated with ALS progression and prognosis. Front Neurol, 2023. 14: p. 1170360.

13. Mauvais-Jarvis, F., et al., Reduced expression of the murine p85alpha subunit of phosphoinositide 3-kinase improves insulin signaling and ameliorates diabetes. J Clin Invest, 2002. 109(1): p. 141–9.

14. Fox, M., H.R. Mott, and D. Owen, Class IA PI3K regulatory subunits: p110-independent roles and structures. Biochem Soc Trans, 2020. 48(4): p. 1397–1417.

15. Li, X., et al., Deregulated Gab2 phosphorylation mediates aberrant AKT and STAT3 signaling upon PIK3R1 loss in ovarian cancer. Nat Commun, 2019. 10(1): p. 716.

16. Tsay, A. and J.C. Wang, The Role of PIK3R1 in Metabolic Function and Insulin Sensitivity. Int J Mol Sci, 2023. 24(16).

17. Sánchez-Alegría, K., et al., PI3K Signaling in Neurons: A Central Node for the Control of Multiple Functions. Int J Mol Sci, 2018. 19(12).

18. Wang, L., et al., Brain Development and Akt Signaling: the Crossroads of Signaling Pathway and Neurodevelopmental Diseases. J Mol Neurosci, 2017. 61(3): p. 379–384.

19. Nogueira, V., et al., Akt determines replicative senescence and oxidative or oncogenic premature senescence and sensitizes cells to oxidative apoptosis. Cancer Cell, 2008. 14(6): p. 458–70.

20. Los, M., et al., Switching Akt: from survival signaling to deadly response. Bioessays, 2009. 31(5): p. 492–5.

21. Du, S. and H. Zheng, Role of FoxO transcription factors in aging and age-related metabolic and neurodegenerative diseases. Cell Biosci, 2021. 11(1): p. 188.

22. Zhang, X., et al., Akt, FoxO and regulation of apoptosis. Biochim Biophys Acta, 2011. 1813(11): p. 1978–86.

23. Winnay, J.N., et al., A regulatory subunit of phosphoinositide 3-kinase increases the nuclear accumulation of X-box-binding protein-1 to modulate the unfolded protein response. Nat Med, 2010. 16(4): p. 438–45.

24. Blanquie, O., et al., Electrical activity controls area-specific expression of neuronal apoptosis in the mouse developing cerebral cortex. Elife, 2017. 6.

25. Lebedeva, J., et al., Inhibition of Cortical Activity and Apoptosis Caused by Ethanol in Neonatal Rats In Vivo. Cereb Cortex, 2017. 27(2): p. 1068–1082.

26. Merling, R.K., et al., Transgene-free iPSCs generated from small volume peripheral blood nonmobilized CD34+ cells. Blood, 2013. 121(14): p. e98–107.

27. Patterson, K., et al., Generation of two tdTomato reporter induced pluripotent stem cell lines (NHLBIi003-A-1 and NHLBIi003-A-2) by AAVS1 safe harbor gene-editing. Stem Cell Res, 2020. 42: p. 101673.

28. Garcia-Montojo, M., et al., TDP-43 proteinopathy in ALS is triggered by loss of ASRGL1 and associated with HML-2 expression. Nat Commun, 2024. 15(1): p. 4163.

29. Wang, T., et al., Derivation of a Human Brain Organoid with Microglia Development. J Vis Exp, 2025(215).

30. Canny, M.D., et al., Inhibition of 53BP1 favors homology-dependent DNA repair and increases CRISPR–Cas9 genome-editing efficiency. Nature Biotechnology, 2018. 36(1): p. 95–102.

31. DeMarino, C., et al., Autophagy Deregulation in HIV-1-Infected Cells Increases Extracellular Vesicle Release and Contributes to TLR3 Activation. Viruses, 2024. 16(4).

