## Supplementary material for "A novel frameshift mutation in Phosphoinositide 3-kinase regulatory subunit 1 (*PIK3R1*) causes immunodeficiency and Amyotrophic Lateral Sclerosis (ALS)": supp figs

### Figure 1 Supp

#### Peripheral Blood

| | Ref. Value | 07/2021 | 02/2017 | 6/2016 | 3/2016 | 1/2016 | 10/2015 | 02/2013 |
|---|---|---|---|---|---|---|---|---|
| **Medication** | | Rituximab discontinued | | | | Rituximab started | | |
| **Blood** | | | | | | | | |
| **CD3#** | 714-2266/mcL | 1196 | 1062 | 1005 | 1236 | 1282 | 889 | 1292 |
| **CD4/CD3#** | 359-1565/mcL | 383 | 392 | 331 | 407 | 291 | 225 | 356 |
| **CD8/CD3#** | 178-853/mcL | 728 | 597 | 603 | 750 | 894 | 577 | 815 |
| **CD19+#** | 61-321/mcL | 30 | 1 | 25 | 0 | 116 | - | - |
| **NK Cell#** | 126-729/mcL | 63 | 35 | 25 | 37 | 65 | 42 | 58 |
| **DNT Cell%** | 1.3-9.2% | | | | | | 8.3 | 7.9 |
| **CD4/CD62L+/CD45RA+/CD3%** | 7.6-37.7% | | | | | | 1.9 | |
| **CD3/CD4/CD62L+/CD45RA-%** | 10.4-30.7% | | | | | | 16.7 | |
| **CD3/CD4/CD62L-/CD45RA-%** | 2.3-15.6% | | | | | | 3.7 | |
| **CD4/CD62L-/CD45RA+/CD3%** | 0.0-1.5% | | | | | | 0 | |
| **CD8/CD62L+/CD45RA+/CD3%** | 5.7-19.7% | | | | | | 20.7 | |
| **CD3/CD8/CD62L+/CD45RA-%** | 1.5-10.3% | | | | | | 17.6 | |
| **CD3/CD8/CD62L+/CD45RA-%** | 1.1-9.2% | | | | | | 10.7 | |
| **CD8/CD62L-/CD45RA+/CD3%** | 0.7-7.8% | | | | | | 8 | |
| **CD20%** | 3.0-19.0% | | | | | | 7.8 | |

#### CSF

| | Ref. Value | 07/2021 | 02/2017 | 6/2016 | 3/2016 | 1/2016 | 10/2015 | 02/2013 |
|---|---|---|---|---|---|---|---|---|
| **Medication** | | Rituximab discontinued | | | | Rituximab started | | |
| **CSF** | | | | | | | | |
| **Glucose** | 40-70 mg/dL | 59 | 50 | 50 | 51 | 49 | 65 | 49 |
| **Protein** | 15.0-40.0 mg/dL | 34 | 42 | 35 | 36 | 29 | 30 | 41 |
| **WBC Count** | | 3 | 11 | 9 | 8 | 7 | 0 | 3 |
| **Lymphocyte %** | | -- | 92 | 94 | 92 | 96 | -- | -- |
| **Oligoclonal Bands** | | 1 CSF, 0 serum | 0 CSF, 0 serum | CSF bands only | 0 CSF, 0 serum | 0 CSF, 0 serum | CSF bands only | CSF bands only |
| **IgG Index** | 0.26-0.62 ratio | 0.33 | 0.4 | 0.35 | 0.55 | 0.45 | 0.54 | 0.58 |

**Figure 2 Supp**

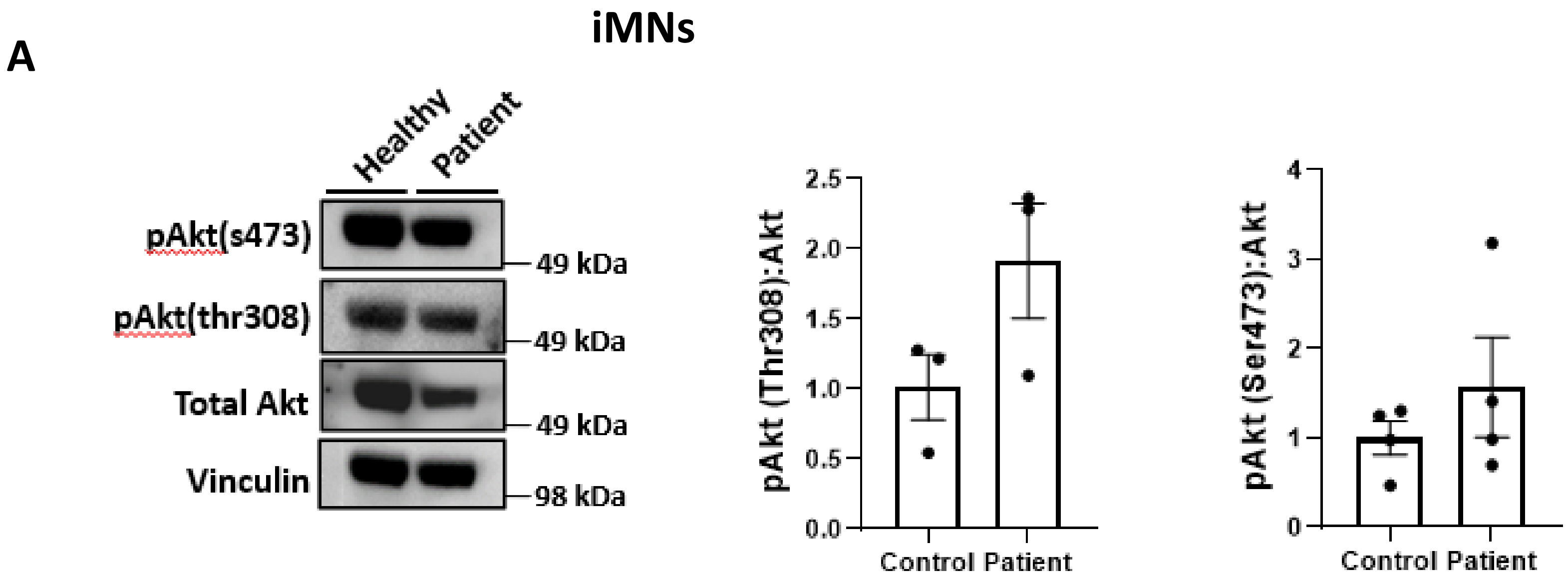


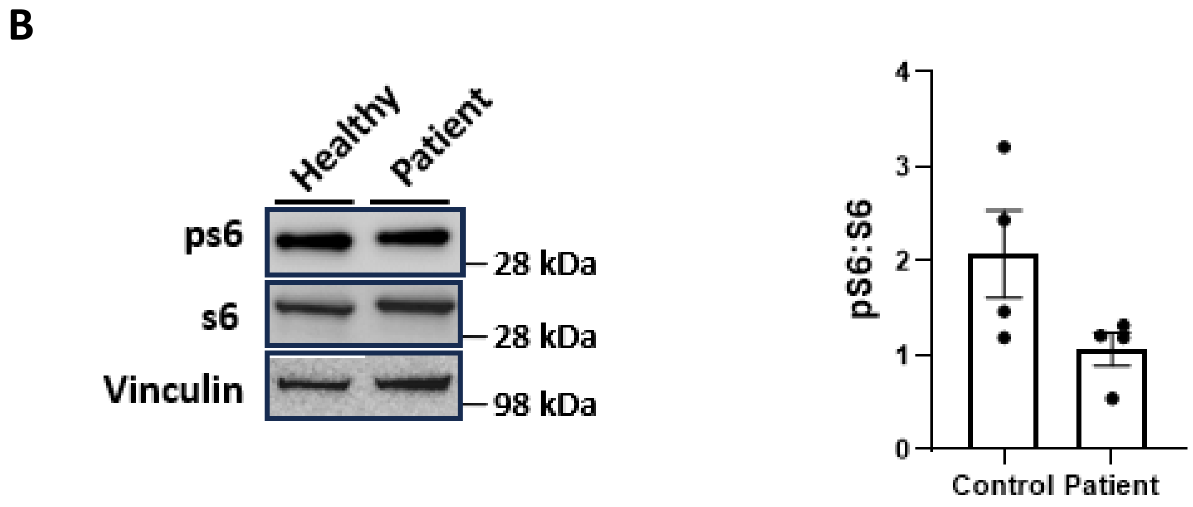

#### Figure 3 Supp

**A**

| | Number of reads | Estimated cell count | Mean reads per cell | Median genes per cell |
|---|---|---|---|---|
| **Mutated** | 811,218,958 | 11,000 | 73,747 | 5,596 |
| **Control** | 878,462,975 | 13,700 | 64,121 | 5,391 |
| **Patient** | 460,223,516 | 4,026 | 114,313 | 2,220 |
| **Healthy** | 1,189,668,728 | 14,785 | 80,465 | 5,790 |

**B**

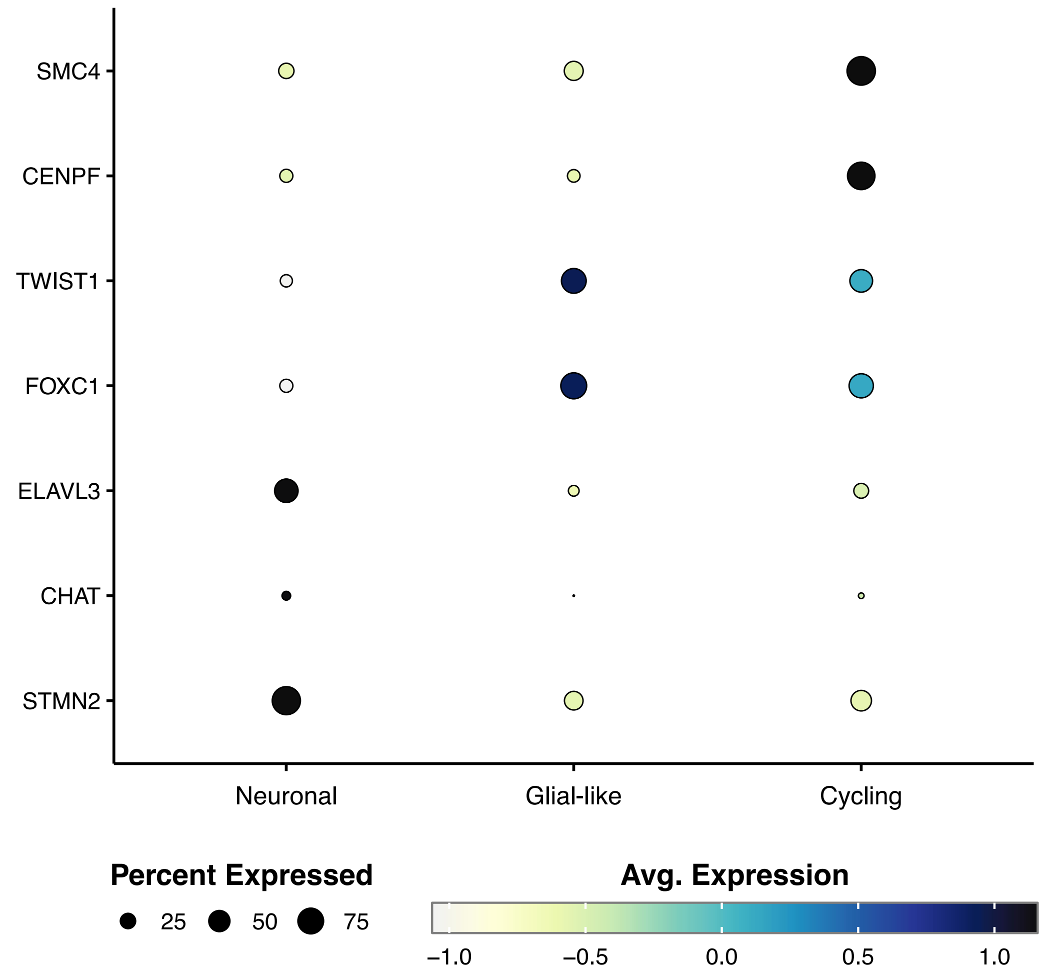


**C**

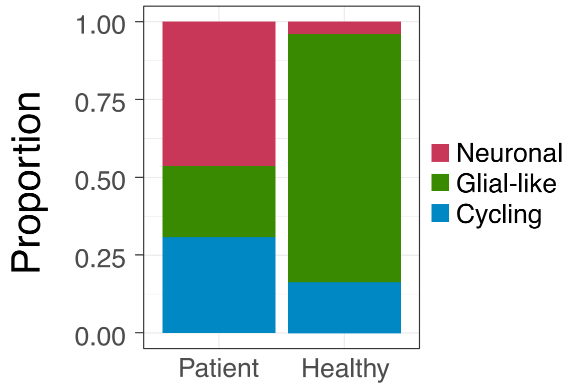

### Figure 4 Supp

**A**

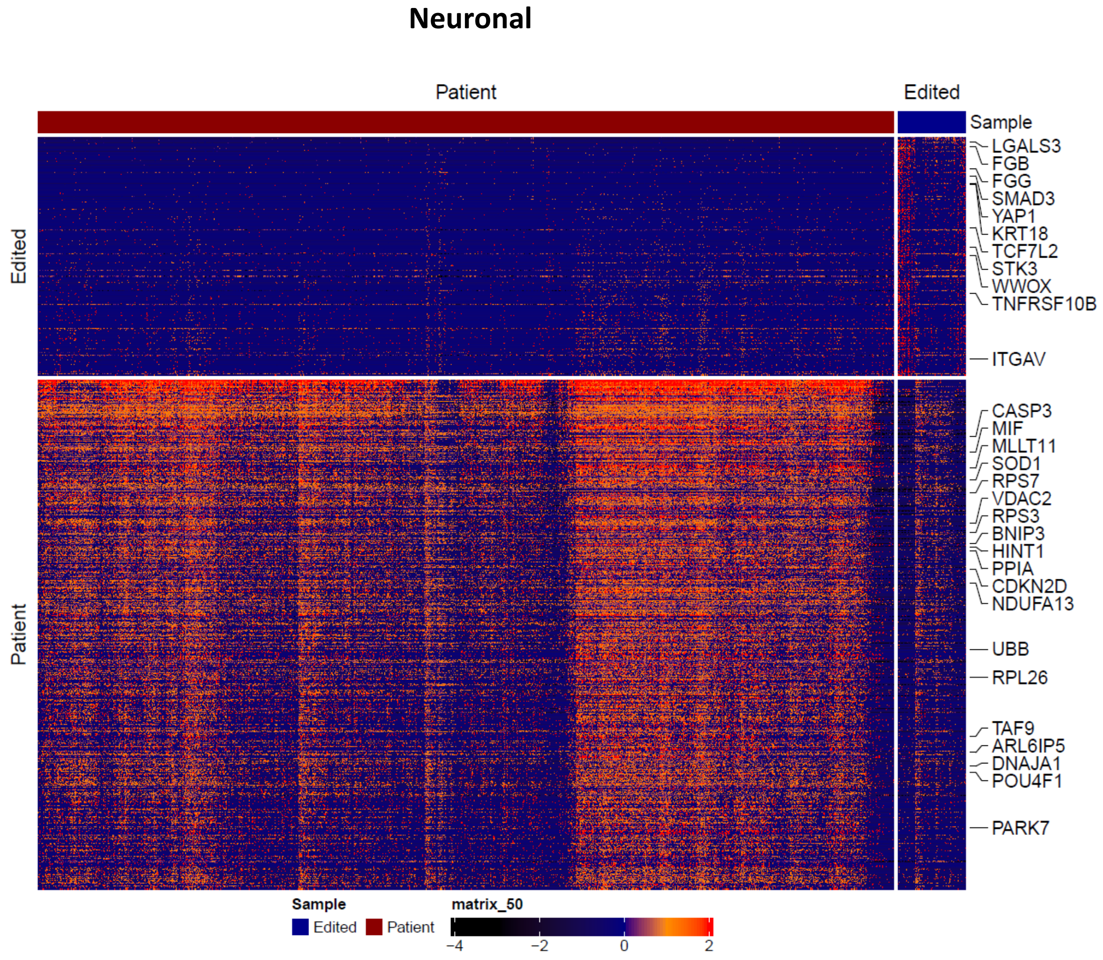


**B**

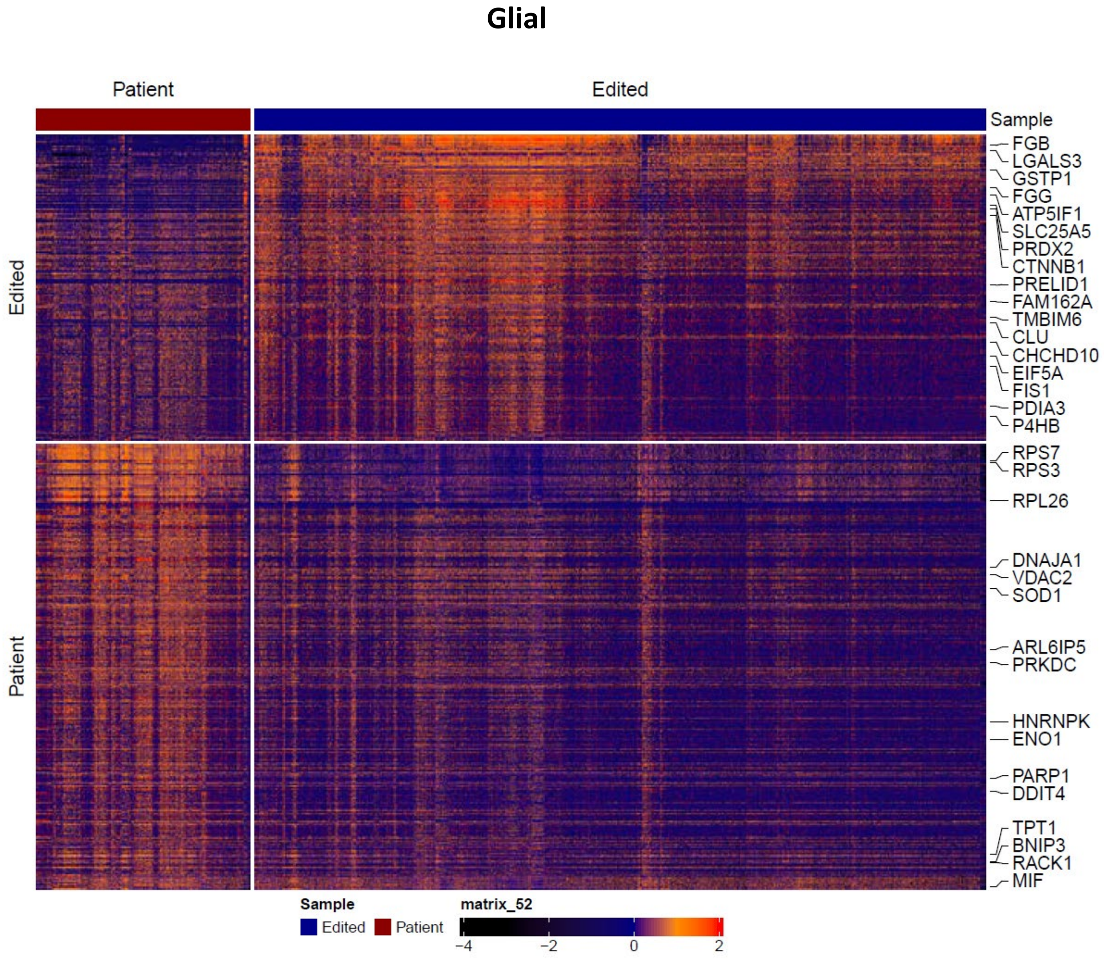

### Table 1 Supp

| Marker | Clone | Fluorophore | Supplier | Cat. # |
|---|---|---|---|---|
| CD45 | 2D1 | PerCP | BioLegend | 368505 |
| CD8 | 3B5 | Qdot 800 | Thermo Scientific | Q22157 |
| CD16 | 3G8 | BUV496 | BD | 612944 |
| CD45RA | 5H9 | BUV395 | BD | 740315 |
| CD123 | 6H6 | Super Bright 436 | Thermo Scientific | 62-1239-42 |
| CD335 | 9E2 | PE | BioLegend | 331908 |
| TCR γδ | B1.1 | PerCP-eFluor 710 | Thermo Scientific | 46-9959-42 |
| CD11c | B-Ly6 | BUV661 | BD | 612967 |
| CD28 | CD28.2 | BV605 | BioLegend | 302968 |
| CD19 | | spark NIR 685 | Biolegend | 302270 |
| CD4 | SK3 | cFlour YG584 | Cytek | R7-20041-100T |
| CD95 | DX2 | PE-Cy5 | BioLegend | 305610 |
| PD-1 | EH12.2H7 | BV421 | BioLegend | 329920 |
| CD86 | FUN-1 | BB515 | BD | 564544 |
| CXCR3 | G025H7 | BV650 | BioLegend | 353730 |
| CCR6 | G034E3 | BV711 | BioLegend | 353436 |
| CCR7 | G043H7 | BV785 | BioLegend | 353230 |
| CD20 | HI47 | Pacific Orange | Thermo Scientific | MHCD2030 |
| CD127 | HIL-7R-M21 | APC-R700 | BD | 565185 |
| CD38 | HIT2 | APC-eFluor 780 | Thermo Scientific | 47-0389-41 |
| CD57 | HNK-1 | FITC | BioLegend | 359604 |
| CD161 | HP-3G10 | eFluor 450 | Thermo Scientific | 48-1619-41 |
| IgD | IA6-2 | BV480 | BD | 566138 |
| CD11b | ICRF44 | PerCP-Cy5.5 | BioLegend | 301328 |
| HLA-DR | L243 | BV570 | BioLegend | 307637 |
| CD25 | M-A251 | PE-Cy7 | BioLegend | 356108 |
| CD24 | ML5 | PE/Dazzle594 | BioLegend | 311134 |
| CD27 | M-T271 | APC | BioLegend | 356410 |
| CD14 | MφP9 | BUV563 | BD | 741441 |
| Viability | N/A | LIVE DEAD Blue | Thermo Scientific | L23105 |
| CD56 | NCAM16.2 | BUV737 | BD | 564447 |
| CD3 | OKT3 | BV510 | BioLegend | 317332 |
| CD33 | P67.6 | Alexa Fluor 647 | BioLegend | 366626 |
| CXCR5 | RF8B2 | BV750 | BD | 747111 |
| CD45RO | UCHL1 | BUV805 | BD | 748367 |
